# The sodium-proton exchangers sNHE and NHE1 control plasma membrane hyperpolarization in mouse sperm

**DOI:** 10.1101/2024.03.04.583310

**Authors:** Analia G Novero, Paulina Torres Rodríguez, José L De la Vega Beltrán, Liz J Schiavi-Ehrenhaus, Guillermina M Luque, Micaela Carruba, Cintia Stival, Iñaki Gentile, Carla Ritagliati, Celia M Santi, Takuya Nishigaki, Diego Krapf, Mariano G Buffone, Alberto Darszon, Claudia L Treviño, Dario Krapf

## Abstract

Sperm capacitation, crucial for fertilization, occurs in the female reproductive tract and can be replicated *in vitro* using a medium rich in bicarbonate, calcium, and albumin. These components trigger the cAMP-PKA signaling cascade, proposed to promote hyperpolarization of the mouse sperm plasma membrane through activation of SLO3 K^+^ channel. Hyperpolarization is a hallmark of capacitation: proper membrane hyperpolarization renders higher *in vitro* fertilizing ability, while *Slo3* KO mice are infertile. However, the precise regulation of SLO3 opening remains elusive. Our study challenges the involvement of PKA in this event and reveals the role of Na^+^/H^+^ exchangers. During capacitation, calcium increase through CatSper channels activates NHE1, while cAMP directly stimulates the sperm-specific NHE, collectively promoting the alkalinization threshold needed for SLO3 opening. Hyperpolarization then feeds back Na^+^/H^+^ activity. Our work is supported by pharmacology, and a plethora of KO mouse models, and proposes a novel pathway leading to hyperpolarization.

**Teaser:** Alkalinization of sperm cytoplasm activates potassium channels to hyperpolarize the plasma membrane in a PKA independent cascade.

## Introduction

The sperm capacitation process in mammalian sperm involves a complex cascade of events that unfolds within the female reproductive tract upon ejaculation (for a review see^1^). These events are vital for enabling sperm to successfully fertilize the egg and are replicated *in vitro* using a specialized capacitating media. These media contain various components, including energy sources, bicarbonate, Ca^2+^, and albumin. At the molecular level, capacitation is hallmarked by critical processes such as sperm membrane potential (*E*m) hyperpolarization and intracellular alkalinization^2–4^. Genetic and pharmacological investigations have unveiled specific transporters that underlie the ionic fluxes involved in such changes. Among them, the atypical sperm-specific Na^+^/H^+^ exchanger (SLC9C1 aka sNHE) and the SLO3 K^+^ channel have emerged as key players.

Their indispensability for capacitation is underscored by the fact that mice deficient in these genes exhibit infertility, albeit without discernible abnormalities in other physiological aspects, as these proteins are uniquely expressed in sperm ^2,5,6^. Given the importance of sperm *E*m hyperpolarization associated to SLO3 currents, it is intriguing that the molecular events leading to SLO3 opening have not been completely revealed yet, besides knowing that SLO3 responds to alkalinization.

The principal membrane H^+^ transporters responsible for mediating H^+^ efflux in response to the rising extracellular pH within the female reproductive tract are Na^+/^H^+^ exchangers (NHEs) (for a review see ^7^). Three groups identify NHE isoforms in mammalian sperm: NHE1 and NHE5 (SLC9A subgroup), NHA1 and NHA2 (SLC9B subgroup), and sNHE as a member of the SLC9C subgroup, with the recent identification of SLC9C2 in at least rat and human sperm^8,9^. Their roles and significance in sperm physiology remain an active area of research.

The localization of NHE1 to the sperm midpiece is established ^10^, but its precise contribution to sperm fertility has been somewhat enigmatic. Notably, the breeding outcomes of NHE1-deficient mice have provided intriguing insights. While mating between *Nhe1* KO males and females yielded no successful breeding, a litter could be carried to term when *Nhe1*^+/-^ (HET) males mated with *Nhe1* KO females ^11^. As for NHE5, its functional importance in sperm physiology is yet to be elucidated, though it has been also localized to the midpiece of mouse sperm ^10^. Disruptions in the NHA exchangers provoked subfertility in single *Nha1* or *Nha2* KO males, whereas double KO males were rendered completely infertile, marked by severely compromised sperm motility. These phenotypes were associated with attenuated cAMP synthesis by soluble adenylyl cyclase (sAC) and reduced expression of the full-length sAC isoform ^12^. Addition of cell-permeable cAMP analogs rescued sperm motility defects, while fertility defects seemed to arise from deficient acrosome reaction ^13^. Regarding sNHE, it is localized to the principal piece of the sperm flagellum. It was found to affect sperm motility, a phenotype amenable to rescue through the addition of cell-permeable cAMP analogs ^6^. The presence of a cyclic nucleotide-binding domain (CNBD) and a putative voltage-sensor motif on sNHE suggests its potential regulation by cyclic nucleotides and changes in *E*m. It has been suggested that cAMP modulates intracellular pH (pHi) by regulating sNHE activity ^14,15^. While these advances have broadened our understanding of key cellular processes that orchestrate capacitation, many questions remain unanswered regarding the precise physiological roles of these transporters (and even their presence) in sperm function.

In this article, we investigate the role of cAMP and its downstream targets in sperm alkalinization that drives hyperpolarization of *E*m during capacitation. Unexpectedly, inhibition of PKA catalytic activity did not impair *E*m hyperpolarization despite sAC activity being necessary for this change in *E*m. Non-capacitated sperm exposed to cAMP permeable analogs underwent *E*m hyperpolarization. Our results using a battery of genetic mouse models and pharmacology demonstrate the critical role of NHE1 and sNHE in the pathway leading to *E*m hyperpolarization, through pH modulation by Ca^2+^ and cAMP, respectively.

## RESULTS

### PKA catalytic activity is not required for *E*m hyperpolarization

One of the initial events in the capacitation-signaling pathway involves the bicarbonate-induced stimulation of sAC, leading to increased cAMP synthesis, which activates PKA ^16^. Two pieces of evidence support PKA’s role in mouse sperm *E*m hyperpolarization: 1) the PKA inhibitor H-89 prevented *E*m hyperpolarization, when present during capacitation, and 2) *E*m hyperpolarization increased with bicarbonate in a concentration-dependent manner ^17^. To gain insights into this pathway, we attempted to inhibit *E*m hyperpolarization using the synthetic and permeable PKA inhibitor peptide sPKI, known to effectively and specifically inhibit its catalytic activity ^18^. Surprisingly, and in contrast to the effect of H-89, sperm *E*m hyperpolarization associated to capacitation was not inhibited by sPKI (Fig. 1A and B). As a control we show that sPKI did not affect *E*m in NC sperm (Fig. 1A and B). We verified the effective blockade of PKA activity by sPKI through immunoblotting analysis using antibodies against phosphorylated PKA substrates (pPKAs) ^19^ (Fig. 1C). Furthermore, *E*m in capacitating medium containing sPKI depended on SLO3 activity, as cells did not hyperpolarize in the presence of BaCl_2_ (Sup. Fig. 1), an effective blocker of SLO3 currents ^20^. As an additional control, sperm from *Slo3* null mice did not hyperpolarize in the presence of sPKI, further disregarding the possibility of non-physiological hyperpolarization caused by sPKI (Fig. 1D).

**FIGURE 1.**
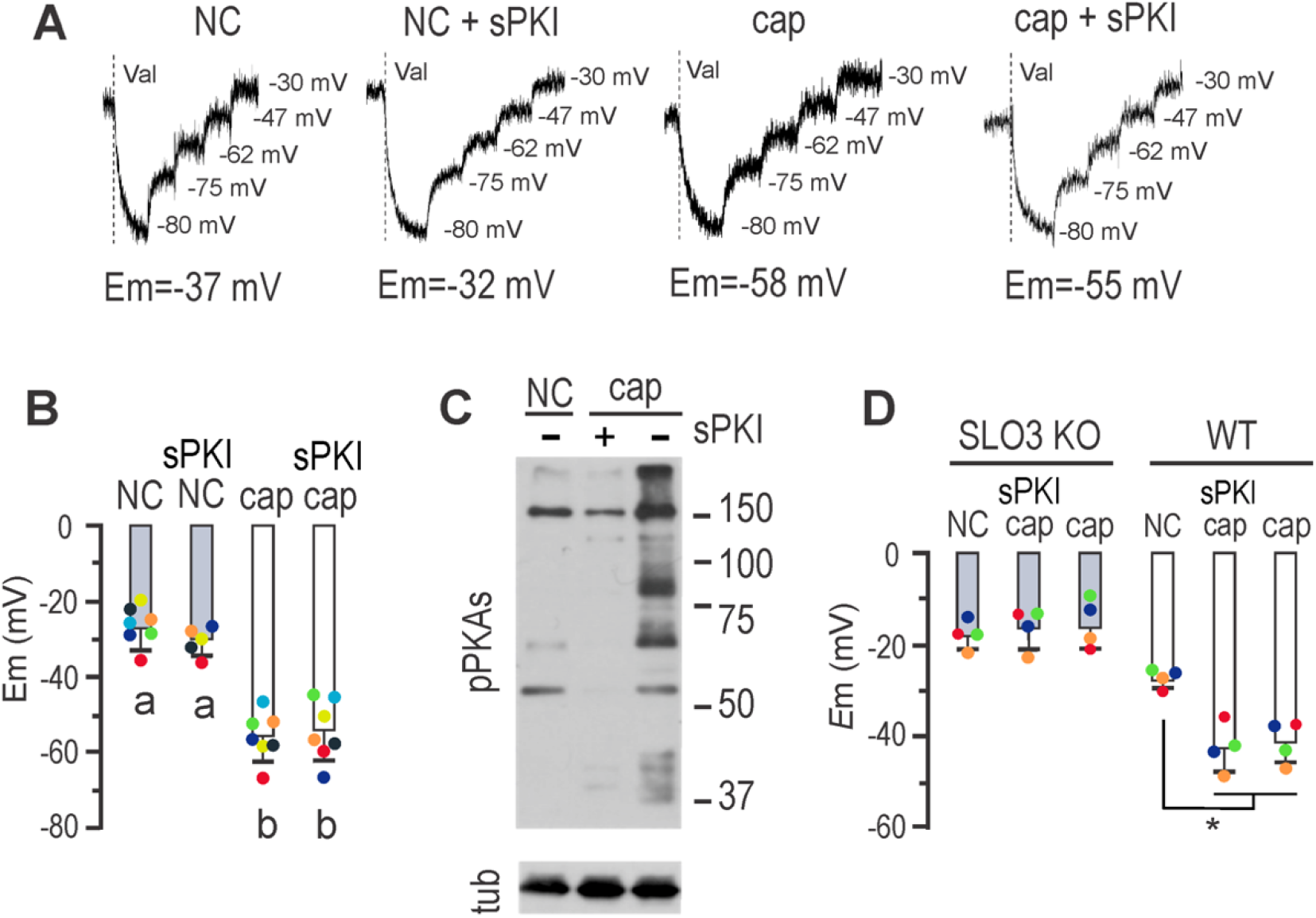
Mouse sperm *E*m hyperpolarization can dispense PKA catalytic activity. ***A***, Fluorescence traces showing the values of the sperm *E*_m_ obtained after sperm incubation in either non-capacitating or capacitating conditions containing or not 15 μM sPKI for 60 min. Each experiment displays its calibration curve and the estimated *E*m value. ***B***, Summary of *E*_m_ measurements of sperm incubated in conditions depicted in *A* (mean ± SEM; *n* ≥ 5); different letters indicate statistically significant differences (*p*<0.001). ***C***, Sperm were incubated for 60 min in non-capacitating or capacitating medium containing or not 15 μM sPKI. Each condition was processed for western blot analysis with a monoclonal anti-pPKAs antibody. Membrane was stripped and analyzed for the presence of tubulin using anti-β-tub (clone E7). ***D***, Sperm *E*_m_ measurements obtained after 60 min incubation of either *Slo3* KO (gray boxes) or WT sperm (white boxes) in either non-capacitating or capacitating medium containing or not 15 μM sPKI (mean ± SEM; *n* = 4; ^*^ *p*<0.001). Each colored dot represents the value for each independent sample.

### cAMP drives *E*m hyperpolarization

Non-capacitating media were supplemented with the permeable cAMP analogue 8Br-cAMP, known to stimulate phosphorylation of PKA substrate ^21^. 8Br-cAMP promoted *E*m hyperpolarization, which sPKI did not inhibit (Fig. 2A). We further inhibited cAMP production during capacitation using the potent and selective sAC inhibitor TDI-10229 ^22^. As shown in figure 2B, TDI-10229 prevented *E*m hyperpolarization when added to capacitating media. The addition of 8Br-cAMP overcame the inhibitory effect of TDI-10229, further supporting the role of cAMP in *E*m hyperpolarization, independently of PKA catalytic activity.

**FIGURE 2.**
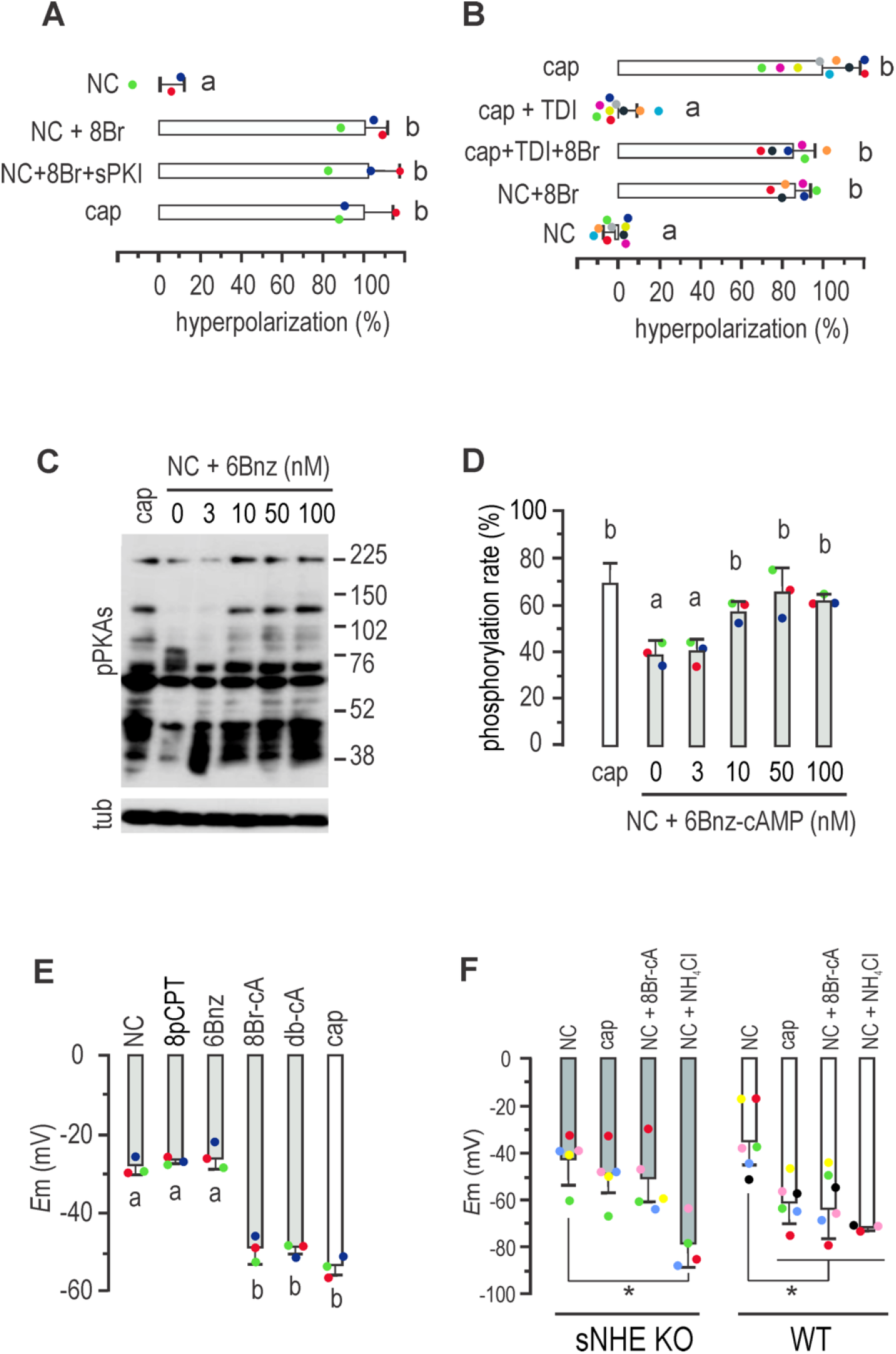
Mouse sperm *E*m hyperpolarization is cAMP regulated and sNHE dependent. ***A***, Sperm *E*_m_ obtained after incubation in either non-capacitating or capacitating conditions containing 500 μM 8Br-cAMP alone or in addition to15 μM sPKI for 60 min. Results are expressed as a normalization of percentage of hyperpolarization considering mean NC and cap values as 0% and 100%, respectively (mean ± SEM; *n*=4). Different letters indicate statistically significant differences (*p*<0.001). ***B***, Sperm *E*_m_ obtained after incubation in either capacitating (with either 10 μM TDI10229 alone or in combination with 500 μM 8Br-cAMP) or non-capacitating conditions containing or not 500 μM 8Br-cAMP for 60 min. Results are expressed as a normalized percentage of hyperpolarization considering NC mean and cap mean values as 0% and 100%, respectively (mean ± SEM; *n*=9). Different letters indicate statistically significant differences (*p*<0.001). Each colored dot represents the value for each independent sample. ***C***, Sperm were incubated for 60 min in non-capacitating media containing increasing concentrations of 6Bnz-cAMP, as indicated. Each condition was processed for western blot analysis with a monoclonal anti-pPKAs antibody. The membrane was stripped and analyzed for the presence of tubulin using anti-β-tub (clone E7). ***D***, Summary of densitometry analysis of sperm cells incubated as in A (mean ± SEM; *n*=3). Different letters indicate statistically significant differences (*p*<0.05). ***E***, *E*_m_ measurements of sperm incubated in non-capacitating conditions containing either 30 μM 8-pCPT-2’-O-Me-cAMP (8pCPT), 50 nM 6Bnz-cAMP, 500 μM 8Br-cAMP or 500 μM dibutyryl-cAMP (db-cAMP) for 60 min (mean ± SEM; *n* = 3). Different letters indicate statistically significant differences (*p*<0.001). ***F***, Summary of *E*_m_ sperm measurements from *sNhe* KO (gray boxes) or WT mice (black boxes) obtained after incubation in either capacitating or non-capacitating conditions containing 500 μM 8Br-cAMP or 10 mM NH_4_Cl for 60 min (mean ± SEM; *n* = 5; ^*^ *p*<0.05).

### cAMP targets during promotion of Em hyperpolarization

The second messenger cAMP has three primary effectors: PKA, the exchange protein activated by cAMP (EPAC, a guanine-nucleotide-exchange factor), and the cyclic nucleotide binding domains found in cyclic nucleotide-gated ion channels (for a comprehensive review, see ^1^). To shed light on the signaling pathway through which cAMP promotes sperm *E*m hyperpolarization, we selected specific agonists of cAMP targets, including the widely used 8-pCPT-2’-O-Me-cAMP to activate EPAC and the membrane-permeant PKA selective agonist N6-Benzyladenosine-cAMP (6Bnz-cAMP). To select an appropriate concentration of 6Bnz-cAMP, we initially exposed sperm to increasing concentrations of the PKA agonist for 60 minutes in non-capacitating media to assess PKA activity, as indicated by the *in vivo* phosphorylation of PKA substrates. Western blots using anti-pPKAs antibodies revealed that a concentration of 50 nM 6Bnz-cAMP induced a saturating phosphorylation of PKA substrates (Figs. 2C and D) and used hereafter to stimulate PKA. Then, the roles of EPAC and PKA in *E*m were then addressed by direct stimulation. As previously shown, 50 μM 8-pCPT-2’-O-Me-cAMP was used for direct stimulation of EPAC^23^. While both general cAMPs analogues db-cAMP and 8Br-cAMP promoted *E*m hyperpolarization, neither specific activation of EPAC with 8-pCPT-2’-O-Me-cAMP nor of PKA with 6Bnz-cAMP induced this *E*m shift (Fig. 2E), ruling out the direct involvement of EPAC and PKA in this process.

The presence of a cyclic nucleotide binding domain in sNHE suggests that its function may be regulated by cAMP. As pharmacological inhibitors of sNHE are not yet available, we analyzed *E*m hyperpolarization in sperm from sNHE null mice. Figure 2F demonstrates that sperm from *sNhe* KO mice did not undergo hyperpolarization when incubated in capacitating conditions for 60 minutes. Gain of function was not observed in the presence of the agonist 8Br-cAMP, which could be attributed to the absence of the cyclic nucleotide binding domain harbored by sNHE. Alkalinization induced by NH_4_Cl addition promoted *E*m hyperpolarization in both WT and sNHE KO sperm, confirming the functional response of SLO3 channels in this KO model.

Interestingly, previous research showed that the sAC inhibitor TDI-10229 blocked pHi increase associated with capacitation ^22^. As mentioned, sNHE is proposed to be also regulated by *E*m through a voltage-sensor domain (VSD), which would be part of a positive feedback loop ^24^. To further support previous work, we analyzed the effect of *E*m on pHi, through the stimulation of *E*m hyperpolarization with the K^+^ ionophore, valinomycin ^25^. Non-capacitated sperm incubated with valinomycin showed an increase of pHi similar to that observed in capacitated cells, as evidence by fluorescence increase of BCECF loaded sperm (Fig. 3A-C), in agreement with previous results ^26^. Of note, this effect on pHi was absent in sNHE deficient mice^26^.

**FIGURE 3.**
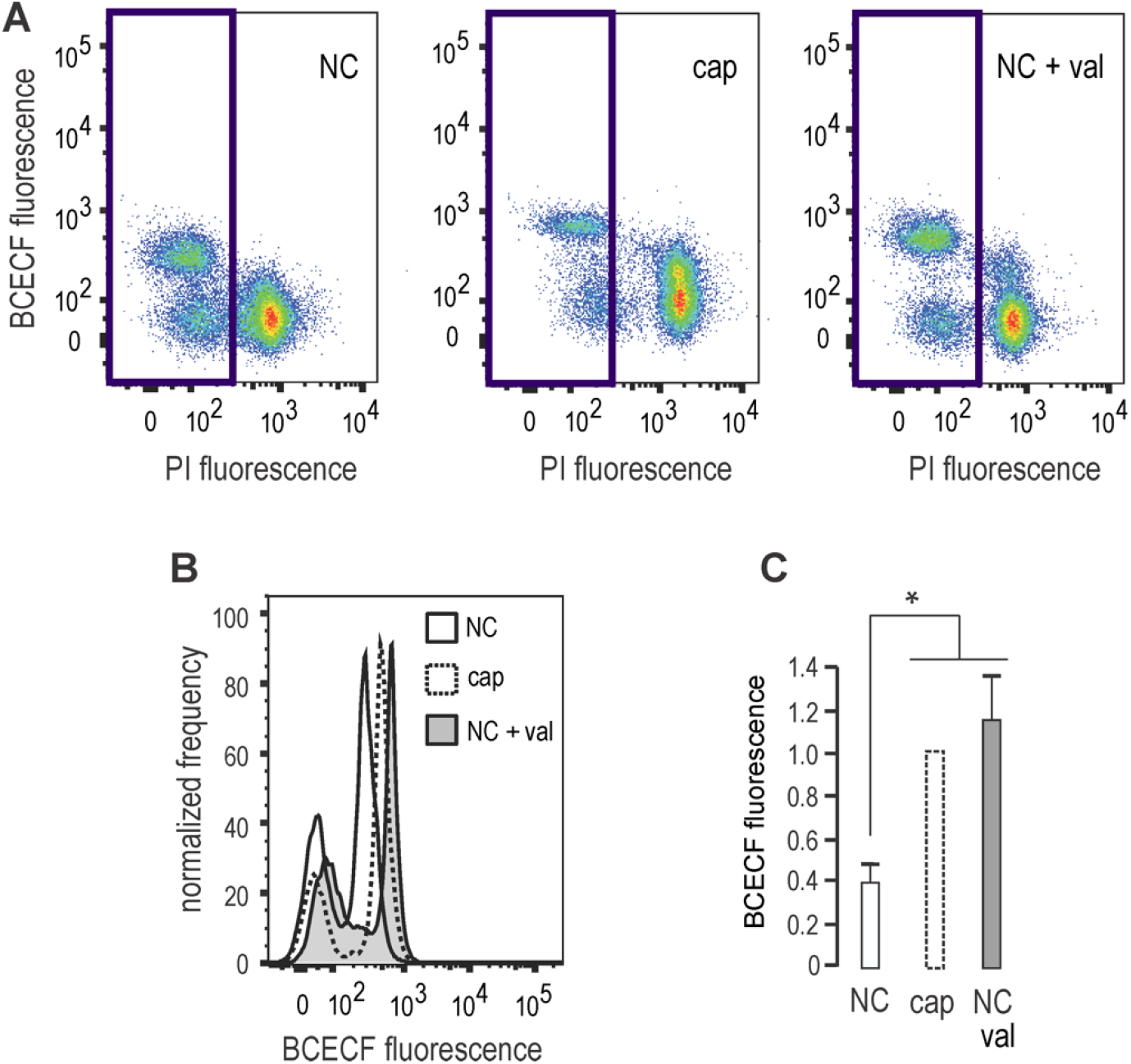
*E*m hyperpolarization induces intracellular alkalinization. Sperm were incubated in either non-capacitating or capacitating media in the absence (DMSO) or presence of 1 μM Valinomycin (Val). ***A***, Representative BCECF versus PI twoLJdimensional fluorescence dot plot analysis. Blue square shows non-PI stained sperm. ***B***, Histogram analysis depicting normalized frequency of sperm, and BCECF fluorescence performed in live sperm populations. ***C***, Normalized median fluorescence intensity of BCECF compared to capacitating condition (mean ± SEM, n=4, ^*^*p* < 0.05).

### Role of Na^+^/H^+^ exchangers in sperm *E*m hyperpolarization

sNHE is not the sole exchanger present in mouse sperm. To explore the role of other Na^+^/H^+^ exchangers in *E*m hyperpolarization, we employed the potent inhibitor 5-(N,N-dimethyl)-amiloride (DMA), which targets members of the SLC9A sub-family, including NHE1, NHE2, and NHE3, with K_i_ values of 0.02, 0.25, and 14 μM, respectively, and negligible effects on NHE4, NHE5, and NHE7 ^27^. DMA, which does not directly affect SLO3 or CatSper channels as recently shown^28^, was added at different concentrations to capacitating media to address its effect on *E*m hyperpolarization. Figure 4A shows a concentration-dependent inhibition of *E*m hyperpolarization. A concentration of 1 μM DMA significantly inhibited hyperpolarization. Of note, 10 μM DMA did not affect the phosphorylation of PKA substrates (Fig. 4B), thus excluding an effect on cAMP synthesis. Considering that NHE1, NHE5, NHA1, NHA2 and sNHE are known to be present in mouse sperm plasma membrane, and that DMA inhibits NHE1, NHE2, and NHE3, then NHE1 might be the primary target of DMA in these cells ^27^. In line with these findings, recent research demonstrated that DMA reduced K^+^ currents in mouse sperm by impairing the regulation of pHi ^28^. Along this line, figure 4C shows that DMA inhibited alkalinization associated to capacitation in mouse sperm.

**FIGURE 4.**
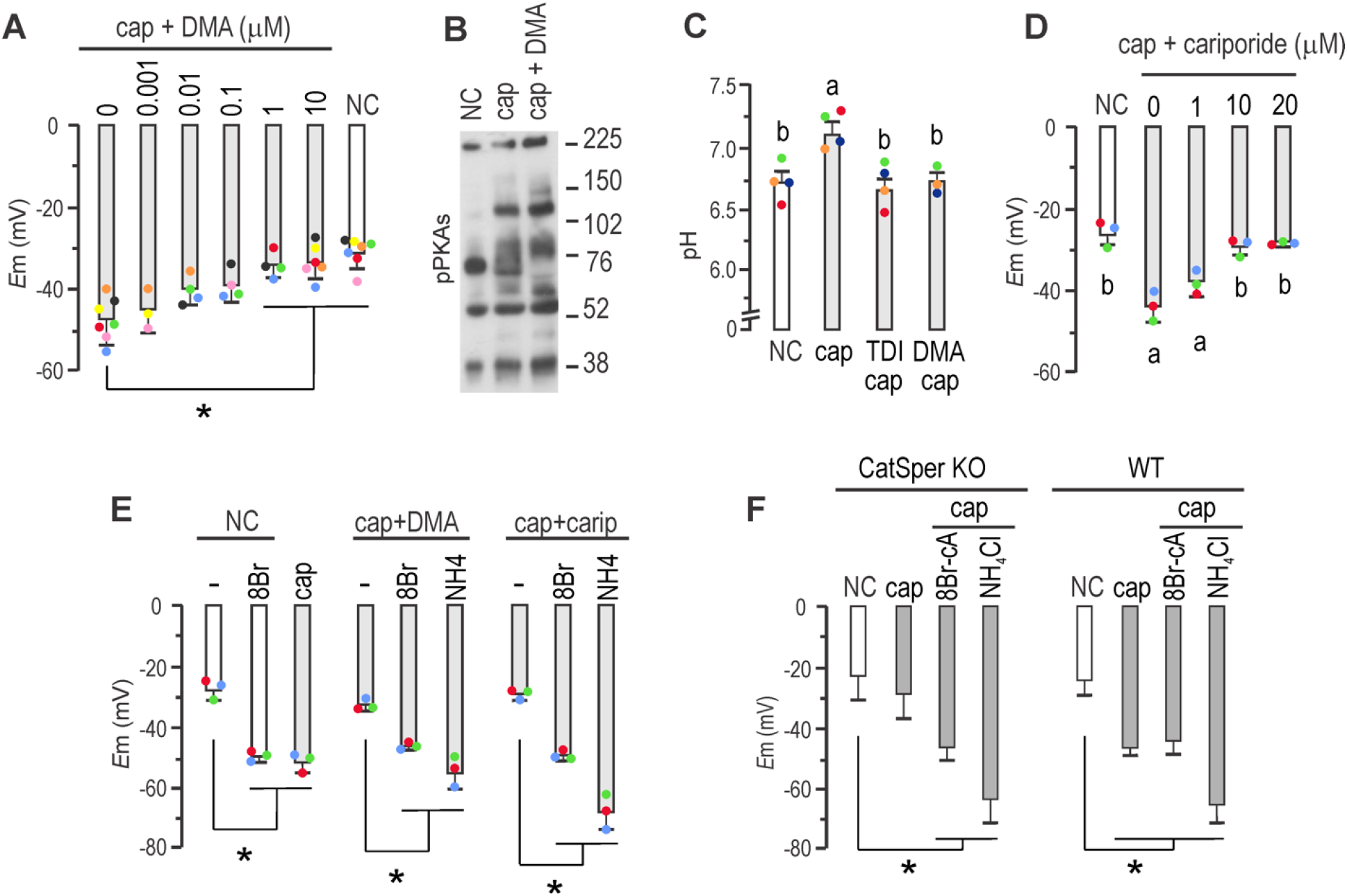
NHE1 activity through Ca^2+^ stimulation is conducive to *E*m hyperpolarization. ***A***, *E*_m_ obtained after sperm incubation in either non-capacitating or capacitating conditions containing different concentrations of dimethyl-amiloride (DMA) as indicated, for 60 min (mean ± SEM; *n* = 4; ^*^ *p*<0.001). ***B***, Sperm were incubated for 60 min in non-capacitating or in capacitating medium in the presence or absence of 10 μM DMA. Each condition was processed for western blot analysis with a monoclonal anti-pPKAs antibody. ***C***, Sperm were incubated for 60 min in non-capacitating or capacitating medium containing or not 10 μM TDI-10229 (TDI) or 1 μM DMA. Different letters indicate statistically significant differences (*p*<0.001). ***D***, *E*_m_ obtained after sperm incubation in either non-capacitating or capacitating conditions containing different concentrations of cariporide, as indicated, for 60 min. Different letters indicate statistically significant differences (mean ± SEM; *n* = 4; ^*^ *p*<0.001). ***E***, *E*_m_ obtained after sperm incubation in non-capacitating conditions in the presence or not of 500 μM 8Br-cAMP (8Br), or in capacitating conditions. As specified, capacitating conditions were supplemented with either 1 μM DMA (DMA), or 10 μM cariporide (carip) for 60 min. As indicated, these conditions were also supplemented with 500 μM 8Br-cAMP or 10 mM NH4Cl (mean ± SEM; *n* = 4; * *p*<0.01). ***F***, Sperm *E*_m_ from either *CatSper1* KO (left panel) or WT mice (right panel) were obtained after sperm incubation in either non-capacitating (with boxes) or capacitating conditions (grey boxes) containing or not either 500 μM 8Br-cAMP or 10 mM NH_4_Cl for 60 min (mean ± SEM; *n* = 5; ^*^ *p*<0.005).

A second inhibitor, cariporide, which selectively targets NHE1 ^29^, was also tested. Figure 4D shows that it effectively inhibited *E*m hyperpolarization when added to capacitating media at 10 μM. The effects of both cariporide and DMA could be bypassed by the addition of either 8Br-cAMP that directly stimulates sNHE or NH_4_Cl that directly stimulates SLO3 (Fig. 4E).

NHE1 possesses a C-terminal domain with extensive disordered regions ^30^, which binds calmodulin in the presence of Ca^2+^^31^. In other cell types, this binding induces an alkaline shift in the pHi sensitivity of NHE1, resulting in its activation at less acidic pHi ^32^. Therefore, the increase in intracellular Ca^2+^ associated with capacitation could activate NHE1, leading to an increase in pHi. To investigate the role of Ca^2+^ in *E*m hyperpolarization, we assessed *E*m in the CatSper1 KO model. Figure 4F demonstrates that sperm from CatSper1 KO mice exhibited deficient *E*m hyperpolarization, which could be restored by the addition of either 8Br-cAMP or NH_4_Cl, similar to the effects observed when inhibiting NHE1 by either DMA or cariporide (see Fig. 4E).

Accordingly, Figure 5 (A-B) shows that the Ca^2+^ ionophore A23187 promoted pHi increase in sperm loaded with the pH sensitive dye BCECF. Treatment with Ca^2+^ ionophore A23187 promoted a pH increase also in sNHE KO, but to a greater extent in sNHE KO than in WT sperm, probably through over-expression of NHE1. As a control, NH_4_Cl was used to evoke pHi increase in both WT and sNHE KO sperm. Same effect was observed when sperm were challenged with ionomycin, as a second Ca^2+^ ionophore (Fig. 5C). Even though Ca^2+^ ionophores are used to increase intracellular Ca^2+^, it can be argued that they also promote pharmacological pHi increase due to H^+^ extrusion, as ionophores exchange Ca^2+^ for H^+^. Thus, as a second approach, cells were incubated in media without added Ca^2+^ salts (which still contain micromolar concentrations of Ca^2+ 33^), and challenged with 1.7 mM Ca^2+^ (Fig. 5D). As before, pHi increase could be clearly evidenced, both in WT and sNHE KO sperm, further substantiating the role of Ca^2+^ on pHi increase and supporting the lack of hyperpolarization in CatSper KO sperm.

**FIGURE 5.**
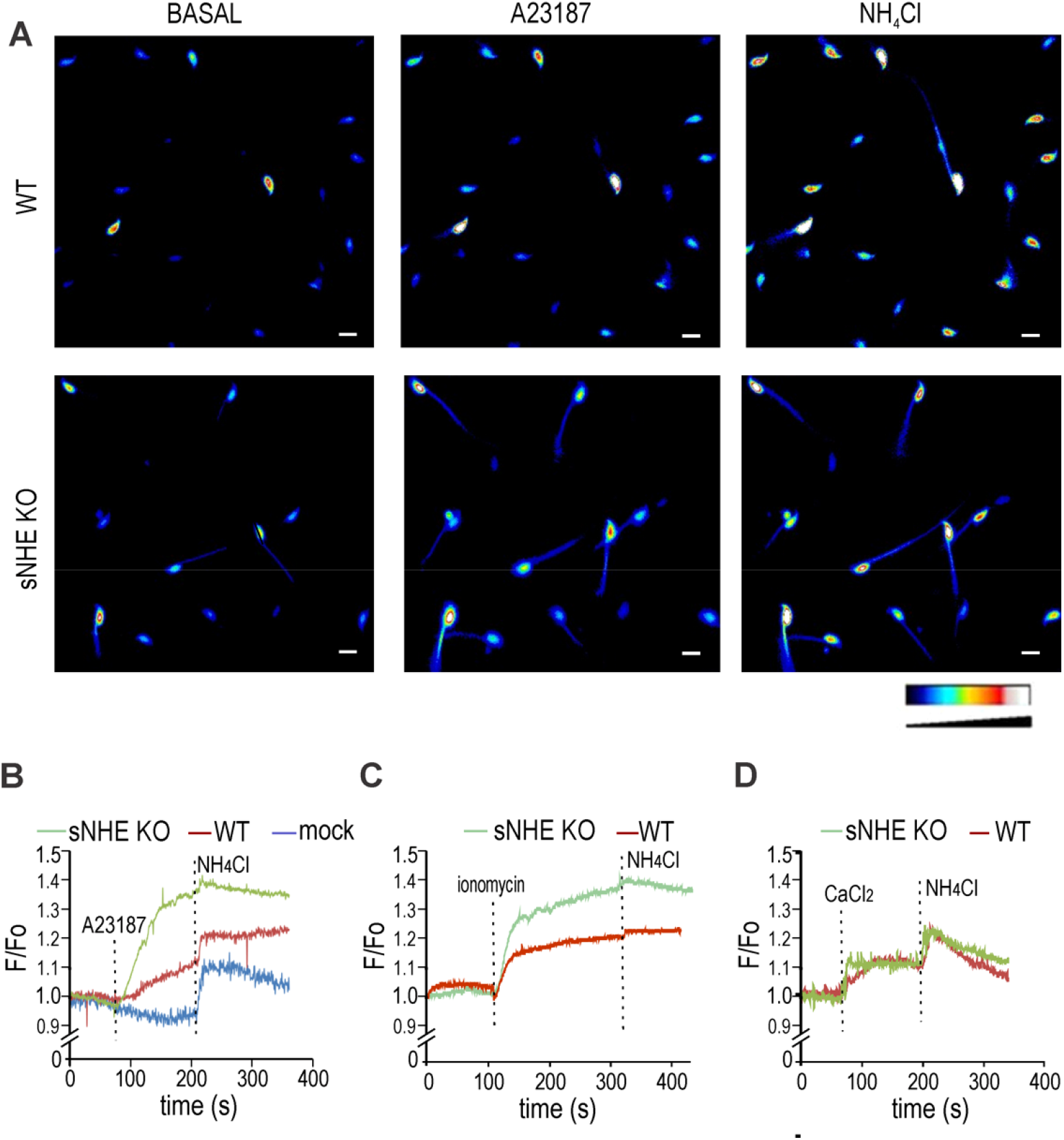
Intracellular Ca^2+^ increase promotes cytoplasmic alkalinization in sperm cells. Non-capacitated sperm cells were loaded with 0.5 μM BCECF-AM for 30 min before smearing onto laminin-precoated coverslips to record fluorescence. ***A***, Representative fluorescence images of WT (upper panels) and *sNhe* KO (lower panels) sperm exposed to 10 μM of the ionophore A23187, followed by 10 mM NH_4_Cl. Reference bar for fluorescence intensity is depicted. Scale bar is equal to 10 μm. ***B***, Summary average traces (8 cells in each trace) of experiments performed in A, including a mock treatment on WT sperm performed with DMSO instead of A23187. ***C***, Summary average traces (8 cells in each trace) of either WT or *sNhe* KO sperm were exposed to 10 μM ionomycin followed by 20 mM NH_4_Cl. ***D***, Summary average traces (8 cells in each trace) of sperm incubated in nominal zero Ca^2+^ (no added Ca^2+^ salts) challenged with 1.7 mM CaCl_2_ and followed by 20 mM NH_4_Cl.

Altogether, these findings indicate that cAMP is responsible for *E*m hyperpolarization, which includes an increase in pH_I_ mediated by NHE exchangers, independent of PKA catalytic activity, and ultimately promoting SLO3 channel opening.

## DISCUSSION

The mammalian sperm-specific K^+^ channel, SLO3, plays a pivotal role in processes leading to sperm capacitation. *Slo3* KO mice are unable to undergo a physiologically stimulated acrosome reaction and are consequently infertile ^2,34^, underscoring the significance of investigating sperm *E*m hyperpolarization. However, the regulation of this channel during capacitation remains poorly understood.

Experiments conducted by Escoffier et al ^35^ demonstrated that *E*m hyperpolarization associated to capacitation was inhibited by H-89. It is worth noting that H-89 is now recognized for its non-specific effects ^36^. On the other hand, synthetic short peptides of PKI, such as PKI-(14-22)-amide (sPKI), have gained wide acceptance as pharmaceutical agents for selectively inhibiting PKA activity ^37^, demonstrating a high level of specificity. Our observations showed that sperm capacitated in the presence of sPKI, while displaying inhibition of PKA substrates phosphorylation, exhibited hyperpolarized *E*m. Furthermore, inhibition of sAC by TDI-10229 ^22^ blocked *E*m hyperpolarization, consistent with its impact on impairing pHi alkalinization ^22^.

When the permeable analogue 8Br-cAMP was introduced to non-capacitated sperm, it induced *E*m hyperpolarization, a finding that aligns with previous reports involving a different cAMP analogue ^35^. This supports the role of cAMP in the road of activating SLO3 channels and excludes PKA as indispensable.

The mouse SLO3 channel was found to be activated by intracellular alkalinization ^38^, though the precise mechanism of its modulation by pHi remains unresolved (for an in-depth review, see ^39^). An important regulator of SLO3 is the leucine-rich-repeat-containing protein 52 (LRRC52) ^40^. The significance of LRRC52 for SLO3 activity was demonstrated in *Lrrc52* KO mice, where alkalinization failed to hyperpolarize sperm *E*m to the same extent as in WT sperm, suggesting a crucial role for LRCC52 in SLO3 response to alkalinization ^39^. In this regard, Na^+^/H^+^ exchangers (NHEs) have emerged as potential contributors to sperm alkalinization during capacitation. NHEs are responsible for regulating the pH of different cell compartments in a variety of cell types. Of particular importance in sperm physiology, sNHE is located in the principal piece of the sperm flagellum and possesses a cyclic nucleotide-binding domain ^41^.

Sperm from sNHE null mice did not undergo capacitation associated hyperactivation ^6^, although this phenotype was restored by the addition of permeable cAMP analogues. Similar to slc9c1 KO, disruption of either NHA1 or NHA2 resulted in a reduced sperm motility phenotype, which was also rescued by incubating sperm with cAMP analogues, pointing towards the role of NHE exchangers in proper regulation of sAC activity and/or expression ^12^. However, our findings herein demonstrate that cAMP analogues did not restore *E*m hyperpolarization in sNHE null sperm. Despite the ability of cAMP addition to restore motility in sNHE null sperm ^6,41^, it did not reinstate *E*m hyperpolarization, indicating the involvement of sNHE in this pathway, likely through its cyclic nucleotide-binding domain. Recently, a human case was reported in which a mutation in sNHE resulted in a deletion in the cyclic nucleotide-binding domain. These sperm, lacking a functional cyclic nucleotide-binding domain, exhibited asthenozoospermia and infertility, indicating its importance also in human sperm ^42^.

Physiological modulation of the NHE family has been a subject of study for many years, mainly through pharmacology. Modified analogues of amiloride were designed to enhance specificity toward NHEs. DMA bears a double substitution of the 5-amino group nitrogen which increases its potency and selectivity toward NHE1. Our results demonstrated that DMA produced a robust inhibition of *E*m hyperpolarization when present in capacitating media. Cariporide, a non-related amiloride inhibitor of NHE1, with negligible effects on either Na^+^/Ca^2+^ exchangers or ENaCs ^43^, also inhibited *E*m hyperpolarization. In all instances, *E*m hyperpolarization could be reinstated by the addition of permeable cAMP analogues. Therefore, these results indicate the

participation of both sNHE and NHE1 in triggering *E*m hyperpolarization of mouse sperm. cAMP possibly acts through the CNBD present in sNHE to increase pHi, triggering the opening of SLO3. This *E*m hyperpolarization, in turn, stimulates a positive feedback loop onto sNHE, as previously suggested ^24,26^. On the other hand, intracellular Ca^2+^ might activate NHE1 through its extensive disordered regions in the long cytoplasmic C-terminal domain ^30,31^. It has been proposed that this binding allows activation of NHE1 at a less acidic pHi ^32^. Therefore, upon the increase in intracellular Ca^2+^ associated with capacitation, NHE1 could drive pHi to a more alkaline state. In human sperm, NHE1 has been recently shown to be expressed at low amounts ^44^. Thus, possible differences in the control of intracellular pH between mouse and human can be expected.

Considering these findings, we propose a mechanism that regulates pHi and *E*m hyperpolarization in mouse sperm, in which NHE1 and sNHE act synergistically (Fig. 6). Synthesis of cAMP induces alkalinization via sNHE, which, together with the action of NHE1 driven by intracellular Ca^2+^ increase, promotes the necessary alkalinization to increase the conductance of SLO3. In turn, *E*m hyperpolarization stimulates a positive feedback loop that further activates sNHE.

**FIGURE 6.**
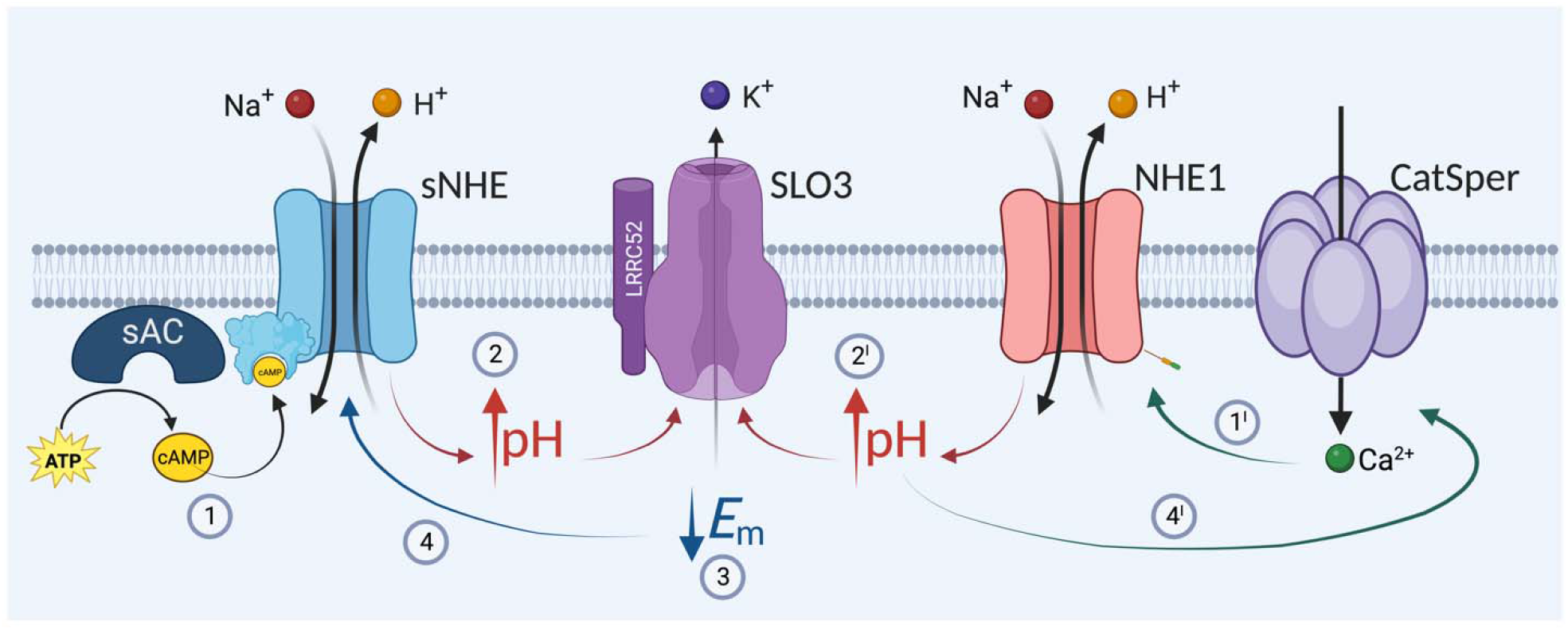
Working model proposing a dual action of sNHE and NHE1 on sperm *E*m hyperpolarization. (1) Synthesis of cAMP induces alkalinization via sNHE, which, together with the action of NHE1 driven by intracellular Ca^2+^ increase (1’s), promote the necessary alkalinization (2 and 2′) to increase the conductance of SLO3 (3). In turn, *E*m hyperpolarization stimulates a positive feedback loop that further activates sNHE (4). It is worth noting that the sole action of sNHE (inhibition of NHE1) or of NHE1 (in the case of *sNhe* KO) is not sufficient, under physiological conditions, to promote SLO3 opening.

This work paves the way for the study of the role of NHEs in mammalian sperm, considering their pivotal role in capacitation. These insights into the regulatory network of sperm capacitation contribute to our understanding of the fundamental processes that underlie fertilization competence, offering new perspectives for future research in this field.

### Materials and Methods Experimental Design

C57BL/6 male mature (10–13 weeks-old) male mice (wild type, *Catsper1* KO ^5^, *Slo3* KO ^2^ and *sNhe* KO ^41^) were used. In all cases, mice housing and all experimental procedures were conducted in accordance with the corresponding Institutional animal care guidelines, reviewed and approved by the Ethical Committees of the Instituto de Biología y Medicina Experimental, Buenos Aires, Argentina #32/2021, Animal Care and Use Committee of the Facultad de Ciencias Bioquímicas y Farmacéuticas de Rosario (UNR), Argentina (#380/2023) and of the Instituto de Biotecnología, UNAM, Mexico. The Guide for Care and Use of Laboratory Animals approved by the National Institutes of Health (NIH) was strictly met. In all cases, sperm were prepared as detailed below, using high grade reagents, as follows: Bovine serum albumin (BSA, fatty acid-free), cariporide (HOE-642), isobutilmetilxantina (IBMX) and 2’-O-dibutiril adenosín monofosfato-3’,5’ cíclico (db-AMPc), carbonyl cyanide 3-chlorophenylhydrazone (CCCP), dimethyl sulfoxide, Ca^2+^ ionophore A23187 and ionomycin were purchased from Sigma (St. Louis, MO). PKI 14–22 amide myristoylated (sPKI) was obtained from Tocris. 3-amino-N-(aminoiminomethyl)-6-chloro-5-(dimethylamino)-2-pyrazinecarboxamide monohydrochloride (DMA), 8-bromo-cyclic 3’,5’-(hydrogen phosphate)-adenosine monosodium salt (8Br-cAMP), (N-benzoyl-adenosine cyclic 3’,5’-(hydrogen phosphate) 6Bnz-cAMP), monosodium salt, and 8-[(4-chlorophenyl)thio]-2’-O-methyl-adenosine cyclic 3’,5’-hydrogen phosphate (8pCPT-2-O’-methyl cAMP), monosodium salt and valinomycin were purchased from Cayman Chemicals (Ann Arbor, MI). Anti-phospho-PKA substrates (anti-pPKAs) (clone 100G7E) antibodies and Horseradish peroxidase-conjugated anti-mouse and anti-rabbit IgG were purchased from Cell Signaling Technology (Danvers,MA). β-tubulin (clone E7) was purchased by Developmental Studies Hybridoma Bank. The sAC inhibitor TDI-10229 was kindly provided by Drs Levin and Buck, Department of Pharmacology, Weill Cornell Medicine, New York City, USA. 3,3-dipropylthiadicarbocyanine iodide (DiSC3(5)), BCECF-AM and pluronic acid from Invitrogen, Thermo Fisher Scientific (Waltham, MA, USA); while propidium iodide from Santa Cruz Biotechnology (Dallas, TX, USA). 3,3-dipropylthiadicarbocyanine iodide (DiSC3(5)), BCECF-AM and pluronic acid were dissolved in DMSO; propidium iodide was dissolved in hexa-distilled water.

### Sperm preparation

Cauda epididymal mouse sperm were collected from adult male mice (10 –13 weeks old). Each minced cauda epididymis was placed in 600 μl of HEPES-buffered TYH medium (H-TYH) containing 119.3 mM NaCl, 4.7 mM KCl, 1.2 mM KH_2_PO_4_, 1.2 mM MgSO_4_, 5.6 mM glucose, 0.5 mM sodium pyruvate, 1.7 mM Ca^2+^, and 20 mM HEPES (pH 7.3), accounting for H-TYH medium (“NC medium”). After 15 min of incubation at 37°C (swim-out), epididymides were removed and the suspension was adjusted with NC medium to a final concentration of 1–2 × 10^7^ cells/ml. For capacitation, BSA and NaHCO_3_ were added to final concentrations of 5 mg/ml and 20 mM respectively (“cap medium”) and incubated at 37°C for at least 1 h, or the indicated period.

### SDS-PAGE and immunoblotting

After treatment, sperm were collected by centrifugation, washed in 1 ml of PBS, resuspended in Laemmli sample buffer without β-mercaptoethanol, and boiled for 5 min. After centrifugation, 5% β-mercaptoethanol was added to the supernatants and boiled again for 5 min. Protein extracts equivalent to 1-2 × 10^6^ sperm per lane were subjected to SDS-PAGE and electro-transferred to PVDF membranes (Bio-Rad) at 250 mA for 60 min on ice. Membranes were blocked with 3% BSA in TBS containing 0.1% Tween-20 (T-TBS). Antibodies were diluted in T-TBS containing 1% BSA as follows: 1/3,000 for anti-pPKAs and 1/10,000 for anti-β-tubulin. Secondary antibodies were diluted 1/10,000 in T-TBS and developed using an enhanced chemiluminescence detection kit (ECL Kallium Biolumina) according to manufacturer’s instructions. When necessary, PVDF membranes were stripped at 60 °C for 15 min in 2% SDS, 0.74% β-mercaptoethanol, and 62.5 mM Tris (pH 6.5) and washed 6 times, 5 min each time, in T-TBS. In all experiments, molecular masses were expressed in kilodaltons (kDa).

### Membrane potential assay in cell populations

Sperm *E*m changes were assessed using DiSC_3_(5), as previously described ^45^. After treatment, cells were loaded with 1 μM of the membrane-potential-sensitive dye DiSC_3_(5) (Molecular Probes) for 2 min. Sperm were transferred to a gently stirred cuvette at 37°C, and the fluorescence was monitored with a Cary Eclipse fluorescence spectrophotometer (Agilent, CA) at 620/670 nm excitation/emission wavelengths. CCCP (0.5 μM) was added as uncoupler of oxidative phosphorylation to avoid mitochondrial contribution to the recorded *E*m. Recordings were initiated when steady-state fluorescence was reached and calibration was performed at the end of each measure by adding 1 μM valinomycin and sequential additions of KCl for internal calibration curves, as previously described ^46^. Sperm *E*m was obtained from the initial fluorescence (measured as Arbitrary Fluorescence Units) by linearly interpolating it in the theoretical *E*m values from the calibration curve against arbitrary fluorescence units of each trace. This internal calibration for each determination compensates for variables that influence the absolute fluorescence values.

### Determination of pHi by flow cytometry

Sperm pHi changes were assessed using BCECF-AM as previously described ^47^. After incubation in the appropriate medium, samples were centrifuged at 400 x *g* for 4 min at room temperature and resuspended in 200 μl of NC H-TYH medium containing 0.5 μM BCECF-AM for 20 min at 37°C. Samples were washed again and resuspended in 50 μl of NC H-TYH medium. Before collecting data, 3 μM of propidium iodide was added to monitor viability. Data were recorded as individual cellular events using a MACSQuant Analyzer cytometer (Miltenyi Biotec, Germany). Side-scatter area (SSC-A) and forward-scatter area (FSC-A) data were collected from 20,000 events per sample in order to define sperm population as previously described ^17^. In all cases, doublet exclusion was performed analyzing two-dimensional dot plot FSC-A vs FSC-H. Positive cells for BCECF were collected using fluorescein isothiocyanate (FITC; 530/30) filter together with peridinin chlorophyll protein complex (PerCP; 670LP) filter. Although the two indicators had minimal emission overlap, compensation was done. For calibration curves, samples were split and resuspended with high K^+^ buffered solutions at pH 6.3, 6.5, 7.0, 7.4, or 8.0 (1.2 mM MgSO4, 1.6mM CaCl_2_, 23.8 mM HEPES, 2.78 mM glucose, 3.38 mM sodium pyruvate, 120 mM KCl; pH previously adjusted with NaOH) and 5 μM nigericin was added to equilibrate intracellular and extracellular pH. Data were analyzed using FlowJo software (V10.0.7).

### Analysis of pHi by single cell imaging

Sperm were loaded with the fluorescent pHi indicator as described in the previous sections. Cells were later adhered to 1 mg/ml laminin-precoated coverslips, allowing their flagella to move continuously. The coverslip was mounted on a chamber (Harvard Apparatus) and placed on the stage of an inverted microscope (Eclipse TE 300; Nikon). Fluorescence illumination was supplied by a Luxeon V Star Lambertian Cyan LED (Lumileds Lighting LLC) attached to a custom-built stroboscopic control box. The LED was mounted into a FlashCube40 assembly with a dichroic mirror (M40-DC400; Rapp Opto Electronic; bandwidths: excitation, 450–490 nm; dichroic mirror 505 nm; and emission, 520–560 nm). The LED output was synchronized to the Exposure Out signal of an iXon 888 CCD camera via the control box to produce a single flash of 2-ms duration per individual exposure. The camera exposure time was set equivalent to the flash duration (2 ms). Images were collected every 500 ms using iQ software (Andor Technology).

### Statistical analysis

Data are expressed as mean ± standard error of the mean (SEM) of at least three independent experiments for all determinations. Statistical analyses were performed using the GraphPad Prism 6 software (La Jolla, CA). Student’s t test was used to compare mean values between control and tested groups, while differences between mean values of multiple groups were analyzed by one-way analysis of variance (ANOVA) with multiple comparison tests. Significance is indicated in the figure legends.

## Acknowledgments

We are thankful to Yoloxochitl Sánchez Guevara for technical support with the *sNhe* KO mouse strain.

## Author contributions

Conceptualization: DarioK, AD, CLT, CMS

Methodology: AN, CS, PTR, GML, JLdlVB, LJS-E, MGB, AD, DiegoK, CLT, TN, DarioK

Investigation: AN, PTR, GML, MC, JLdlVB, LJS-E Visualization: AN, PTR, GML, JLdlVB Supervision: DarioK, Diego K, AD, CLT Writing—original draft: DarioK

Writing—review & editing: All authors reviewed the results, edited the manuscript and approved its final version.

## Data and materials availability

All data are available in the main text or the supplementary materials.

