## Supplementary material for "The sodium-proton exchangers sNHE and NHE1 control plasma membrane hyperpolarization in mouse sperm": Sup. Fig. 1

**SUPPLEMENTARY INFORMATION**

**Supplementary Figure 1.**

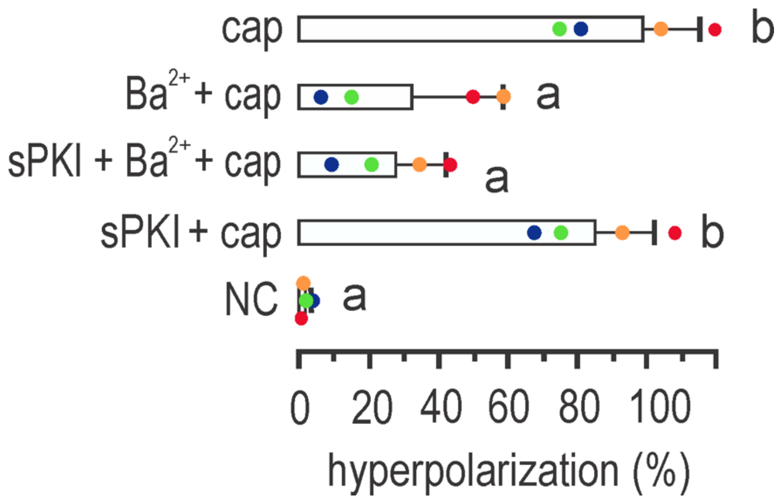


**Legend**

**Supplementary Figure 1. $Ba^{2+}$ blocks *E*m hyperpolarization even in the presence of sPKI.** Sperm $E_m$ measurements obtained after incubation for 60 min in capacitating conditions containing either 1 mM $BaCl_2$ or 15 µM sPKI or a combination of both. Results are expressed as a normalization of percentage of hyperpolarization considering mean NC and cap values as 0% and 100%, respectively (mean ± SEM; $n$=4); different letters indicate statistically significant differences ($p$<0.05).
